# Loss of the Y chromosome in bladder cancer drives metabolic reprogramming

**DOI:** 10.1101/2025.08.26.672471

**Authors:** Boopathi Subramaniyan, Bipul R. Acharya, Prince Anand, Saravana Kumar Kailasam Mani, Saswat Mohapatra, Xingyu Chen, Fangyuan Qu, Yesai Fstkchyan, Corazon Gutierrez, Hany Abdel-Hafiz, Jim Costello, Dan Theodorescu

## Abstract

Metabolic reprogramming promotes cancer aggressiveness and an immune-suppressive tumor microenvironment. Loss of the Y chromosome (LOY) drives both phenotypes in bladder cancer (BC). We investigated the hypothesis that LOY leads to metabolic reprogramming using untargeted metabolomic profiling of human BC cells and analysis of pan-cancer transcriptomic datasets. This revealed that aerobic glycolysis is activated in LOY BC cells. Since prior work showed that expression of collagen receptor DDR2 drives BC progression and DDR2 is a regulator of tumor metabolism, we investigated if DDR2 is implicated in metabolic reprogramming of LOY-tumors. Analysis of scRNAseq data from 251 patients with 12 tumor types found that LOY and DDR2 expression promote aerobic glycolysis, and this was confirmed by metabolomics. Deletion of DDR2 in LOY BC cells reduced glycolytic flux, inhibited cell proliferation, reduced EMT and stemness features, and promoted apoptosis. Our data provide a rationale for using LOY as a tumor selection biomarker for DDR2 targeted therapeutics.

**Teaser:** Loss of the Y chromosome augments glycolytic metabolism in bladder cancer cells, and this is in part dependent on the collagen receptor DDR2.

## Introduction

Loss of the Y chromosome (LOY) in either cancer cells or blood (*1–4*) is associated with aggressive behavior of male cancer. In some tumor types such as bladder cancer (BC) this has shown to be via the creation of an immune-suppressive tumor microenvironment which also makes these tumors more sensitive to immune checkpoint inhibitors (*2*). While the mechanisms that tumor cells with LOY use to drive either aggressive behavior or immune evasion remain to be fully defined, metabolic reprogramming may potentially explain how cells drive both phenotypes (*5, 6*).

A hallmark of cancer, metabolic reprogramming (*7*) allows tumor cells to adjust their energy production and biosynthetic pathways (*6*) to support proliferation, maintain stem-like properties, and evade immune responses (*8–12*). Traditionally the metabolism of cancer cells has been associated with a shift to aerobic glycolysis and lactate production even in the presence of oxygen (*12*). Metabolic reprogramming also upregulates biosynthetic and bioenergetic pathways, such as amino acid metabolism, glutaminolysis, lipid metabolism, mitochondrial metabolism, nucleotide metabolism, and the urea cycle (*13–19*).

In BC the degree of aggressiveness is associated with aerobic glycolysis (*20*). Acidosis of the tumor microenvironment by lactate release hinders immune cell functioning and weakens the effect of immune checkpoint inhibitors (*21–23*). Here, we assess the hypothesis that LOY impacts tumor metabolism in using MB49 murine BC cells (*24*), which have naturally lost the Y-chromosome as well as CRISPR engineered variants of these cells (*2*). We found that LOY cells have a unique metabolite and gene expression profiles known to support tumorigenic growth, survival, and immune escape. Since in our prior work we have shown that expression of the collagen receptor DDR2 determines BC aggressiveness and tumor response to immune checkpoint inhibitors (*25*) is known to influence cancer cell metabolism, particularly glycolysis (*26, 27*), we sought to determine if DDR2 had any impact on the metabolic changes observed in LOY cells. This led us to find that DDR2 depletion can suppress the pro-oncogenic metabolic rewiring in LOY BC cells growth, proliferation, stemness and immune evasion phenotype, and promote cell death. Here we show that LOY is a driver of metabolic reprogramming and that inhibition of DDR2 may reverse these changes. This lays the translational foundation for using LOY as a patient selection biomarker for DDR2 targeted therapeutics that are available clinically.

## Results

### LOY is associated with metabolic reprogramming in bladder cancer

To investigate the metabolic landscape resulting from the loss of Y chromosome (LOY), we used two models of MB49 murine BC cells (*24*). The first model consisted of a pair of MB49 cell lines, one with an intact Y chromosome (YposN) and the other that had naturally lost the Y chromosome (YnegN). These cells were generated by pooling multiple single clones that were found Ypos and Yneg after limiting dilution of MB49 cells henceforth called YposN and YnegN (*2*). A second model was generated by CRISPR engineered deletion of Y chromosome genes in the YposN MB49 cell pool (CRISPR-LOY), henceforth called YposC and YnegC (*2*). Cells were subjected to untargeted ultrahigh performance liquid chromatography–mass spectrometry (UHPLC-MS)-based metabolomics profiling (**Fig. 1A**). A total of in 177 distinct metabolites were detected in YposN and YnegN cells, which were spatially distant in distinct clusters in unsupervised principal component analysis (PCA), indicating changes in YnegN metabolome upon LOY (**Fig. 1B**). The variable importance in projection (VIP) score identified and ranked metabolites that significantly contribute to the differences between groups (**Fig. S1A**). This variation was further supported by the Pearson correlation analysis demonstrating significant negative correlations between these cell lines (**Fig. 1C**). From univariate analysis we identified 84 <u>d</u>ifferentially (absolute fold-change>1.2, adjP<0.1) <u>a</u>bundant <u>m</u>etabolites (DAMs) of which 59 increased and 25 decreased in abundance in YnegN compared with YposN cells (**Fig. 1D** and **S1B**). The hierarchical clustering of DAMs between YnegN and YposN cells showed metabolites either increased or decreased in LOY cells (**Fig. 1E**; top50 DAMs based on fold change, **Fig. S1C**; all metabolites). Further, analysis of metabolic pathways (**Fig. 1F**) and Metabolite Set Enrichment Analysis (MSEA) (**Fig. 1G**) identified Warburg effect (aerobic glycolysis), Purine metabolism, and Arginine and Proline metabolism as highly enriched and impacted metabolic pathways in YnegN cells.

**Figure 1:**
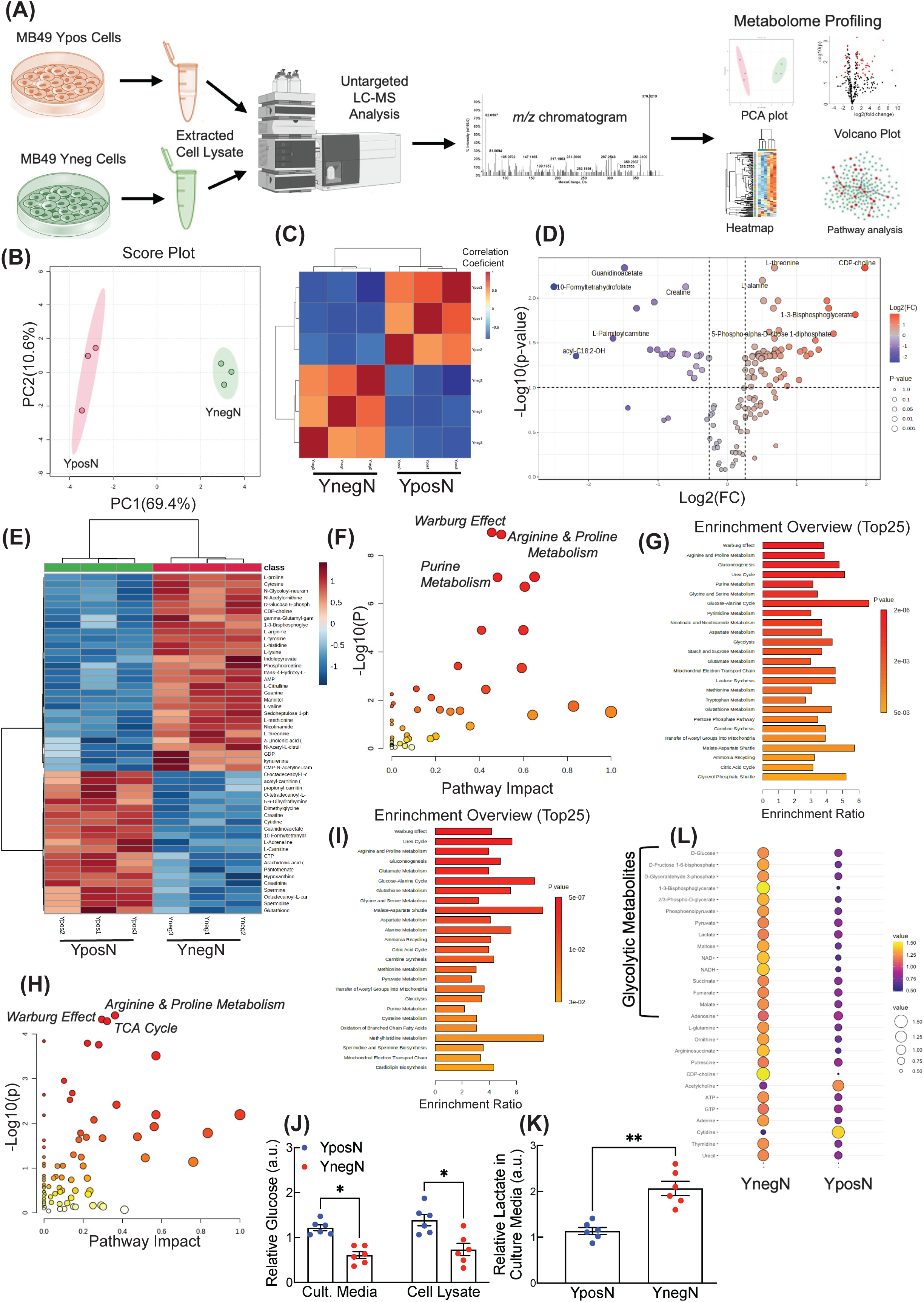
Metabolic Reprogramming in LOY tumor cells. (A) Schema of metabolomics experiments and analysis. (B) PCA analysis showing variance between YposN and YnegN cell metabolome. (C) Pearson Correlation heatmap of variance between YposN and YnegN cells. (D) Volcano plot analysis for significant metabolites. with p-value cut off at 0.1 and fold difference cut off at 1.2. (E) Heatmap of 50 metabolites with highest fold differences in DMA of YnegN vs YposN. (F and G) Metabolic Pathways enrichment analysis (F) and MESA (G) with DMA of YnegN vs. YposN cells. (H and I) Metabolic Pathways enrichment analysis (H) and MESA (I) with DMA of YnegC vs. YposC cells. (J and K) Relative Glucose (J) and Lactate (K) abundance in YposN and YnegN cells. (L) Dot plot showing the relative abundance of Glycolytic- and oncometabolites obtained from significant DMA of YposN vs. YnegN cells. Significance was calculated using pairwise comparison (t-test) with welch correction (K) or with 2-way ANOVA with multiple comparisons (J): *p < 0.05, **p < 0.01.

Similarly, in CRISPR-LOY, spatially distinct clusters of YposC and YnegC samples in unsupervised principal component analysis (PCA) identified metabolite variations among YposC and YnegC cells (**Fig. S1D)**. A total of 155 distinct metabolites were detected and univariate analysis identified 63 DAMs (37 increased and 26 decreased abundance) in YnegC LOY cells (**Fig. S1E**,**F**). The hierarchical clustering of DAMs between YnegC and YposC cells showed metabolites either increased or decreased in LOY cells (**Fig. S1G**; top50 DAMs based on fold change). Metabolic pathway analysis (**Fig. 1H**) and Metabolite Enrichment Set Analysis (MESA) (**Fig. 1I**) identified similar metabolic pathways like Warburg effect (aerobic glycolysis), as being enriched upon LOY in both CRISPR LOY and Natural LOY cell lines. To validate this observation, we checked glucose and lactate levels in cells. In LOY cells, we found reduction in glucose concentration both intracellularly as well as in their cell culture media suggesting glucose uptake from culture media and successive consumption for aerobic glycolysis (**Fig 1J**) (*28*). Similarly, increased lactate abundance was observed in LOY cell lysate (**Fig. 1K**) indicating activation of aerobic glycolysis (*29*). LOY cells also had enrichment in other metabolic pathways such as TCA cycle (**Fig. 1G, H**), arginine and proline amino acid metabolism (**Fig. 1F, H)**, purine metabolism, and the urea cycle (**Fig. 1F**). In addition, several glycolytic metabolites and oncometabolites (succinate, fumarate, arginosuccinate etc.) which regulate cancer cell proliferation, stemness and immune evasion were found enriched in LOY cells (**Fig. 1L**). Taken together, we found that LOY BC cells possess a distinct metabolic landscape that is favorable for their growth, survival, and immune evasion.

### Pathway-enrichment analysis of LOY tumor cells reveals features of stemness and EMT

To gain more insight of LOY induced metabolic reprograming, we performed an integrated-multiomics pathway-enrichment analysis combining differentially expressed genes (DEGs) from transcriptomics and differentially abundant metabolites (DAMs) from metabolomics of LOY vs CONTROL (hencefoth CON; with Y chromosome) MB49 cells (*30*). This integrated analysis identified several metabolic pathways impacted in LOY cells, including glycolysis, gluconeogenesis, purine metabolism, and amino acid metabolism (arginine, proline, etc.) (**Fig. 2A**). Genes encode enzymes to catalyze metabolic reactions. Therefore, differential gene expression in cells can directly affect the production and activity of metabolic enzymes and impact entire metabolic pathways, leading to altered metabolite abundances. Also, metabolites act as regulators of gene expression, creating feedback loops where changes in metabolite levels influence the expression of genes involved in their own production (*31, 32*). We identified differential correlation patterns between DEGs (Total-RNAseq data; YnegC vs. YposC; obtained from *Hany, Nature, 2023* (*2*)) and DAMs in YposN and YnegN cells, suggesting gene expression in these cells influenced their metabolic heterogeneity (**Fig S2A**). Interestingly, correlation between glycolysis genes expression and abundance of glycolytic metabolites was markedly distinct in YnegN and YposN cells (**Fig. 2B**). Using a longitudinal Pan-Cancer single cell transcriptomic dataset comprised of transcriptomic profiles from 497,055 single cells from 346 samples across 251 individuals, collected from 20 publicly available scRNA-seq studies (*33*), we observed higher glycolysis score, a numerical representation of the extent to which glycolysis is active in a cell or tissue (*34*), and higher mean expression of glycolytic genes in LOY cancer cell types (**Fig. 2C,D** and **Fig. S2B,C**).

**Figure 2:**
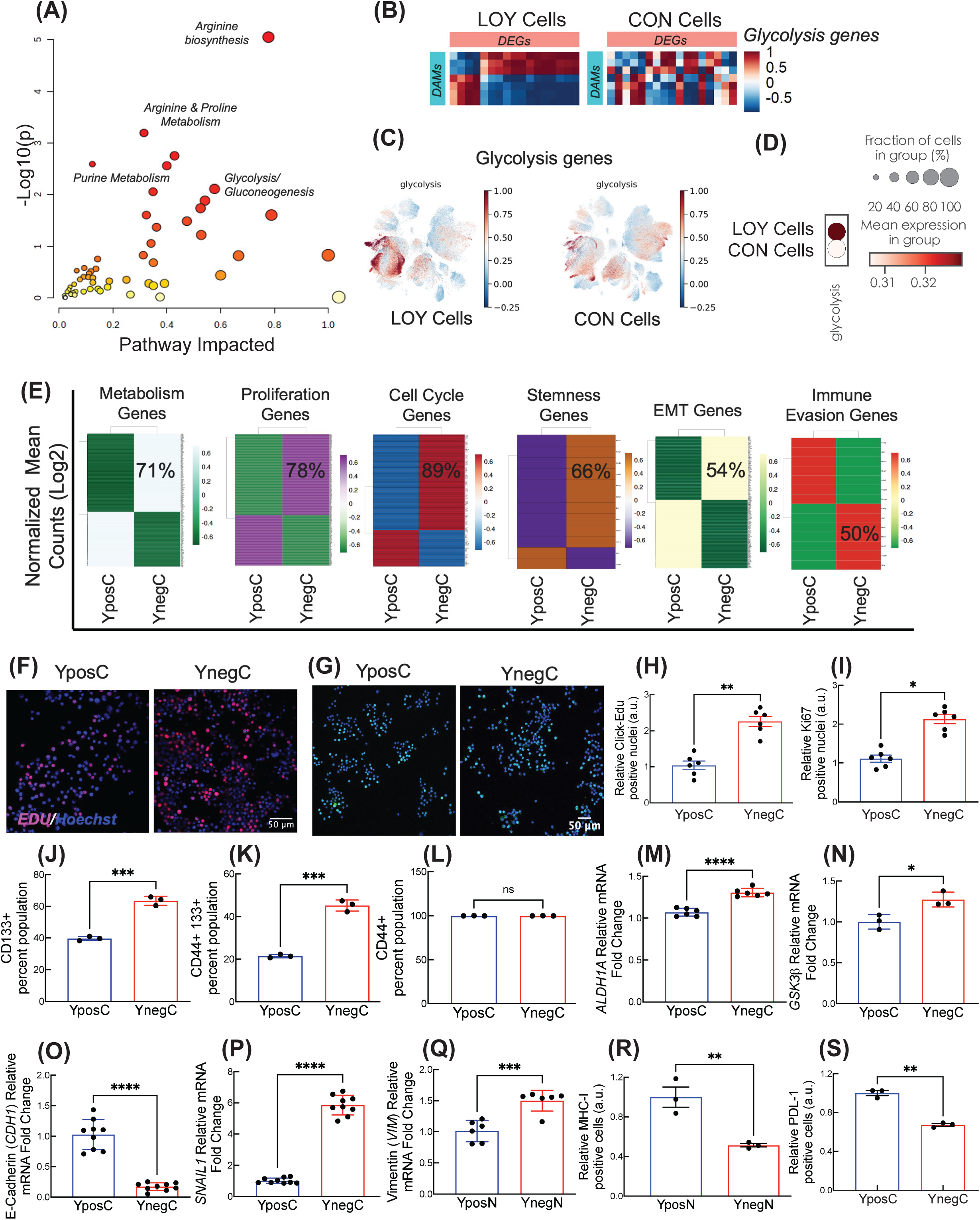
Integrated metabolomics and transcriptomics analysis. (A) Integrated pathways analysis of differential metabolome and transcriptome in YnegN vs. YposN cells. (B) Correlation heatmap showing the DMA and DEG correlation between LOY and CON cells for KEGG glycolysis pathway, the LOY clustering is applied on CON cells, so the metabolites and genes on rows and column are same, to show the differential correlation. (C) Uniform Manifold Approximation and Projection (UMAP) visualization of 497,055 single cells from human tissues profiled by single-cell RNA sequencing, differential UMAP for cells with Y chromosomes (CON Cells) and Cells with loss of Y-chromosome (LOY Cells) is colored by glycolysis gene set enrichment score (GSEA-derived or AddModuleScore), highlighting cell type–specific variation in glycolytic activity. Warmer colors indicate higher glycolytic activity. (D) Dot plot summarizes average glycolytic scores and the proportion of cells expressing glycolysis-related genes across experimental groups (LOY Cells vs. Con Cells). Dot size corresponds to the fraction of cells in each group expressing the pathway, and color intensity reflects the mean module score. Color scale ranges from −0.4 to +0.4. (E) Normalized gene expression analysis from total-RNAseq data of both YposC and YnegC cells, genes belong to the KEGG metabolism genes, proliferation genes, cell cycle genes, and Hallmark geneset of cancer stemness, EMT, and immune evasion genes. (F and H) Click_IT-Edu cell proliferation analysis in YposN and YnegN cells, immunofluorescence images (F) and quantitation (H). (G and I) Ki67 cell proliferation/division analysis in YposC and YnegC cells, images (G) and quantitation (I). (J-L) Cancer stemness surface expression flow cytometry analysis of CD133 (J), CD133/CD44 double positive (K), and CD44 (L) in YposC and YnegC cells. (M-N) Cancer stemness gene expression analysis by qPCR of *ALDH1A* (M) and *GSK3b* (N). (O-Q) EMT gene expression analysis by qPCR of E-cadherin (*CDH1*) (O), *SNAIL1* (P), and Vimentin (*VIM*) (Q). (R and S) Surface expression of PDL-1 (P) and MHC-I (Q) in Ypos and Yneg cells analyzed by flow cytometry. Scale bars: 50 mM (F, G). Significance was calculated using either pairwise comparison (t-test) with welch correction (H-S): *p < 0.05, **p < 0.01, ***p < 0.001, and ****p < 0.0001.

To check if aerobic glycolysis (*35*) in LOY cells regulate cell proliferation, cell survival, cell cycle progression, cancer stemness, EMT, and antitumor immune escape, we profiled the expression status of the genes responsible in LOY cells. Specific pre-annotated genesets for these process were obtained from MSigDB for GO-term, KEGG, of HALLMARK pathways. Normalized transcript counts exhibited higher expression of gene associate with metabolic, proliferation, cell cycle, and stemness conditions in YnegC cells (**Fig. 2E** and **S2D**). We validated these findings with Click-iT EdU assays in LOY cells and determined enhanced cell proliferation in YnegN than YposN cells (**Fig. 2F,H**). Higher Ki-67 index in YnegC cell nuclei further indicated their aggressive growth and continued cell division (**Fig. 2G,I**). As activation of the stemness program in tumor cells ensures their self-renewal (*11*), we investigated the status of stemness markers in LOY cells by quantitative flow cytometric analysis. Higher sub-populations of CD133 as well as CD44+CD133+ (double positive) cells were detected in LOY cells as compared to the Y+ cells (**Fig. 2J,K**), while CD44+ cells remained same in both conditions (**Fig. 2L**). This indicates CD133+ cancer stem cell sub-population enrichment in LOY cells that predominantly influences their self-renewal. Cancer stemness and epithelial–mesenchymal transition (EMT) was further evaluated by quantitative PCR analysis of stemness and EMT marker genes. We observed higher gene expression of cancer stem cell (CSC) marker *ALDH1A* and *GSK3b* in LOY (**Fig. 2M,N**). A reduced expression of E-cadherin (*CDH1*) (**Fig. 2O**) and upregulation of EMT genes *SNAIL1* and Vimentin (*VIM*) was detected (**Fig. 2P,Q**), which indicates that LOY cells have more prominent EMT phenotypic features. Finally, since LOY cells display immune escape, we assessed the expression of MHC-I by flow cytometric analysis and found that MHC-I expression on LOY cell surface was significant reduced in compassion to Y+ cells (**Fig. 2S**), which is an important mechanism of tumor immune evasion, as it prevents cytotoxic CD8+ T cells from recognizing and destroying the cancer cells by reducing the presentation of tumor-associated antigens (*36*). However, in YnegN cells, decreased PD-L1 expression was observed (**Fig. 2R**), suggesting that this loss may hinder anti-PDL1 immunotherapy against aggressive bladder cancers. Our data suggest that metabolic reprogramming in LOY cancer cells provides them with growth and proliferative advantages while reducing their vulnerability to antitumor immune responses.

### DDR2 contributes to the LOY induced reprogramming seen in tumor cells

The collagen receptor DDR2 is a modulator of cancer cells metabolism, particularly of aerobic glycolysis (*26, 27*). Previously, we showed that DDR2 drives bladder tumor aggressiveness and response immune checkpoint therapy (*25*). Hence, we sought to determine if DDR2 had any role in the metabolic changes observed as a function of LOY. We first evaluated the relationship between DDR2 and LOY by examining DDR2 expression levels in TCGA cohort of male bladder cancer patients stratified for high and low expression of Y-chromosome genes, and found that DDR2 expression was higher in Y-Low patients compared to the Y-High cohort (**Fig. 3A**). This result is replicated in our YnegN cell line, which showed higher expression of DDR2 compared to its counterpart; YposN (**Fig. S3A**). We then examined the spatial expression of DDR2 protein in CON (with Y chromosome) and LOY cells in both MB49-LOY models and found that LOY cells have higher DDR2 protein expression in both cytoplasm and nucleus (**Fig. 3B,C** and **S3B,C**), and this was associated with increased phosphorylation (**Fig. S3D**). To evaluate the effect of loss of DDR2 has on LOY tumor metabolism, we generated YnegN DDR2 knockout cell line with CRISPR mediated DDR2 deletion, both immunoblot and immunofluorescence images confirmed the loss of DDR2 in this cell line (**Fig. S3E**). UHPLC-MS-based metabolomics of LOY cells with DDR2 (CON_YnegN) and DDR2KO LOY (DDR2KO_YnegN) cells showed two spatially distinct data clusters on unsupervised principal component analysis (PCA) suggesting metabolome variance between these cell types (**Fig. S3F**). Pearson correlation analysis revealed significant negative correlations between them (**Fig. S3G**). A total of 182 distinct metabolites identified in DDR2KO_YnegN and CON_YnegN cells, which were then subjected to univariate analysis (absolute fold-change>1.2, adjP<0.1), which determined 41 metabolites either increased (n=23) and decreased (n=18) in abundance in DDR2KO_YnegN compared with CON_YnegN cells (**Fig. 3D** and **S3H**). The hierarchical clustering of DAMs shows differential abundance status of metabolites among both the cells (**Fig. 3E**; only 41 DAMs**, Fig. S3I**; all 183 metabolites). In both metabolic pathway enrichment and MESA, the predominant metabolic pathways found activated in DDR2KO_YnegN cells were unrelated to glycolysis, (**Fig. 3F,G**). In contrast, integrated transcriptomic and metabolomic pathway analysis (**Fig. 3H**) identified glucose pathway enrichment in DDR2KO_YnegN cells. By comparing differential expression of genes from the KEGG glycolysis/gluconeogenesis pathway between YnegC vs. YposC and CON_YnegN vs. DDR2KO_YnegN, we found several glycolytic enzymes, such as Hexokinase, Pgam1, Pgk1, and Enolase upregulated in YnegC cells, while these were downregulated in DDR2KO_YnegN cells. This suggests that inhibiting DDR2 in LOY cells leads to a shift away from aerobic glycolysis. (**Fig. S3I**). We further investigated the correlation between DDR2 and glycolytic metabolism in the Pan-Cancer scRNA-seq dataset, described in **fig. 2C** (*33*). Our analysis found a negative correlation between expression of glycolysis related genes in LOY tumor cells without DDR2, with higher expression of glycolytic genes found in LOY tumor cells with DDR2 (**Fig. 3I**). Next, we evaluated the glucose and lactate levels in these cell pairs and found the intracellular glucose level was augmented in DDR2KO_YnegN cells, but media glucose levels were unchanged suggesting the glucose consumption was reduced in the absence of DDR2 (**Fig. 3J**). The lactic acid production was reduced by DDR2 inhibition in DDR2KO_YnegN cells (**Fig. 3K**).

**Figure 3:**
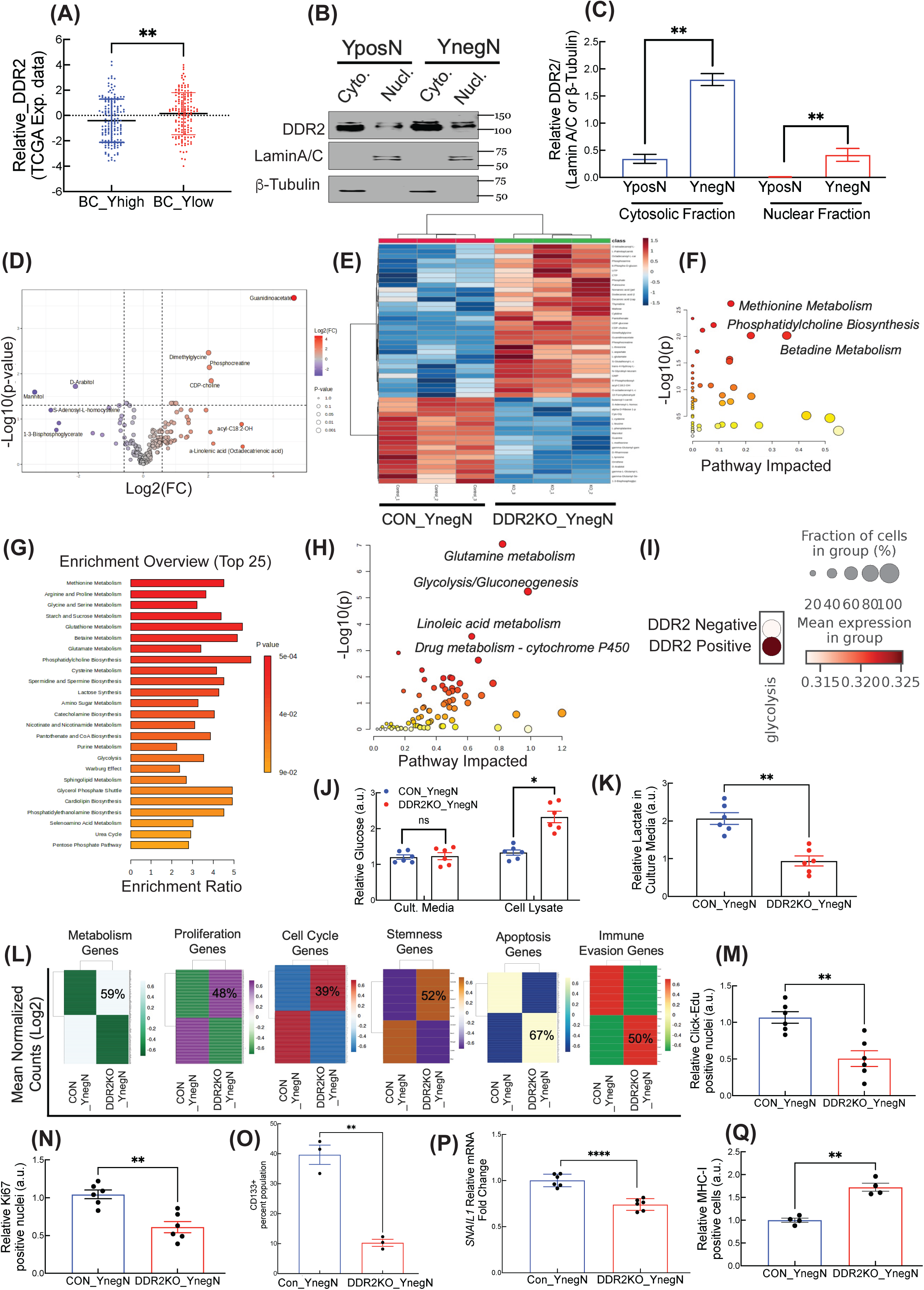
DDR2 modulates metabolic reprogramming in LOY cells. (A) TCGA cohort analysis of gene expressions; DDR2 expression is higher in male BC patients with low Y chromosome gene expression. (B and C) DDR2 protein expression is higher in YnegN cell cytoplasm and nuclei. (D) Heatmap of 50 metabolites with highest fold differences in DMA of DDR2KO_YnegN vs CON_YnegN cells. (E) Volcano plot analysis for significant metabolites. with p-value cut off at 0.1 and fold difference cut off at 1.2. (F and G) Metabolic Pathways enrichment analysis (F) and MESA (G). (H) Integrated pathways analysis of DMA and DEG in DDR2KO_YnegN vs CON_YnegN cells. (I) Dot plot summarizes average glycolytic scores and the proportion of cells, indicating glycolysis-related genes expression across experimental groups; DDR2_Negative and DDR2_Positive). Dot size corresponds to the fraction of cells in each group expressing the pathway, and color intensity reflects the mean module score. Color scale ranges from −0.4 to +0.4. (J and K) Relative Glucose (J) and Lactate (K) abundances in CON_YnegN and DDR2KO_YnegN cells. (L) Normalized gene expression analysis from total-RNAseq data of both CON_YnegN and DDR2KO_YnegN cells, genes belong to the KEGG metabolism genes, proliferation genes, cell cycle genes, and Hallmark geneset of cancer stemness, EMT, and immune evasion genes. (M and N) Quantification of nuclei positive for Click_IT-Edu cell proliferation (M) and Ki67 proliferation/division assay (N). (O) Cell surface expression of cancer stem cell marker CD133 by flow cytometry analysis. (P) *SNAIL1*; EMT gene expression analysis by qPCR (Q) Surface expression of MHC-I on CON_YnegN and DDR2KO_YnegN cells analyzed by flow cytometry. Significance was calculated using either pairwise comparison (t-test) with welch correction (K, M-Q) or with 2-way ANOVA with multiple comparisons (J): *p < 0.05, **p < 0.01, and ****p < 0.0001.

DDR2 deletion also impacted the abundance of metabolites involved in arginine/proline and other amino acids metabolism, glutathione metabolism, nucleotide metabolism, and carnitine and fatty acid metabolism pathways (**Fig. 3D** and **S3I**) involved in cancer (*37*). Transcriptomic profiling of DDR2KO_YnegN cells showed that DDR2 inhibition also inhibited proliferation and cell cycle gene expression while it increased the expression of apoptotic genes (**Fig. 3L**). MESA analysis indicated that the metabolite abundance of both nucleotide metabolism and urea cycle went down indicating DDR2 deletion affected the proliferation of LOY cells (**Fig. 3G**). Reduction in the number of both Edu (5-ethynyl-2’-deoxyuridine) and Ki67 positive nuclei in DDR2KO_YnegN cells compared to its counterpart (**Fig. 3M,S3K** and **Fig. 3N,S3L**) further confirmed DDR2 loss affected proliferation in LOY cells. Flow cytometry analysis showed reduced mean fluorescence intensity of Ki67 in DDR2KO_YnegN cells compared to CON_YnegN, suggesting reduced proliferation **(Fig. S3M).** Depletion of DDR2 in YnegN tumor cells also affected their stemness, as determined by the reduction in the stemness marker CD133+ expressing cell population (**Fig. 3O**), while no reduction was observed in CD44+ cell population, but the mean fluorescence intensity and relative gene expression of CD44 were significantly reduced in DDR2KO_YnegN cells compared to its counterpart (**Fig. S3O-P**). Loss of DDR2 also affected the EMT potential of YnegN tumor cells, as both Snail (**Fig. 3P**) and Vimentin expressions (**Fig. S3R**) were reduced and E-cadherin expression was increased (**Fig. S3Q**). Reduced abundance of oncometabolites like ornithine, kynurenine, and serine, which reduced the immune vulnerability of tumor cells (*38, 39*) was noticed in in DDR2KO_YnegN cells (**Fig. S3I**). DDR2 inhibition also increased surface expression of MHC-I in LOY cells (**Fig. 3Q**) and further reduced the PD-L1 surface expression, supporting the observation that DDR2 depletion enhances antitumor immunity (*25*) (**Fig. S3Q**).

In DDR2KO_YnegN cells, we identified the reduction of HIF-1α (**Fig. 4A**) that directly impacts aerobic glycolysis (*40*) in cancer cells to rapidly produce biomass and energy needed for tumor proliferation and survival. And, as expected, loss of DDR2 in LOY cells impacted their growth, clonogenicity and survivability. We found that DDR2 depletion in LOY cells arrested LOY cell cycle progression at G2/M phase (**Fig. 4B** and **S4A**), loss of survivability was supported by reduced clonogenicity (**Fig. S4B,C**) and loss of viability (in MTS assay, **Fig.4C** and **S4D**). To define the mechanism of loss of cell viability, we identified almost 25-fold abundance of Guanidinoacetate (GAA), a metabolic intermediate primarily synthesized from arginine and glycine (**Fig. 3D**). Abnormal accumulation of GAA has been linked to cellular stress and apoptosis in various cell types, including cancer. GAA disrupts mitochondrial function by increasing reactive oxygen specie (*41*) production that causes oxidative stress, damaging DNA, proteins, and lipids, leading to mitochondrial permeability transition pore (mPTP) opening and cytochrome c release, key steps in apoptosis (*42*). Of note, as determined in Annexin V/7-AAD assay, DDR2KO_YnegN cells were significantly more apoptotic (Early; 37%, late; 9.10%) than CON_YnegN cells (Early; 11.1%, late; 4.14%) (**Fig. 4D** and **S4E**). Loss of DDR2 activated apoptotic signaling hierarchy in LOY cells as it reduced the phosphorylation of both Erk and Src-Kinases, activated cytochrome C and cleaved caspases (**Fig. 4E**). GSEA analysis on DEG list of DDR2KO_YnegN vs. CON_YnegN further validated apoptosis by identifying higher enrichment score of apoptotic genes in DDR2KO_YnegN cells **(Fig. S4F).**

**Figure 4:**
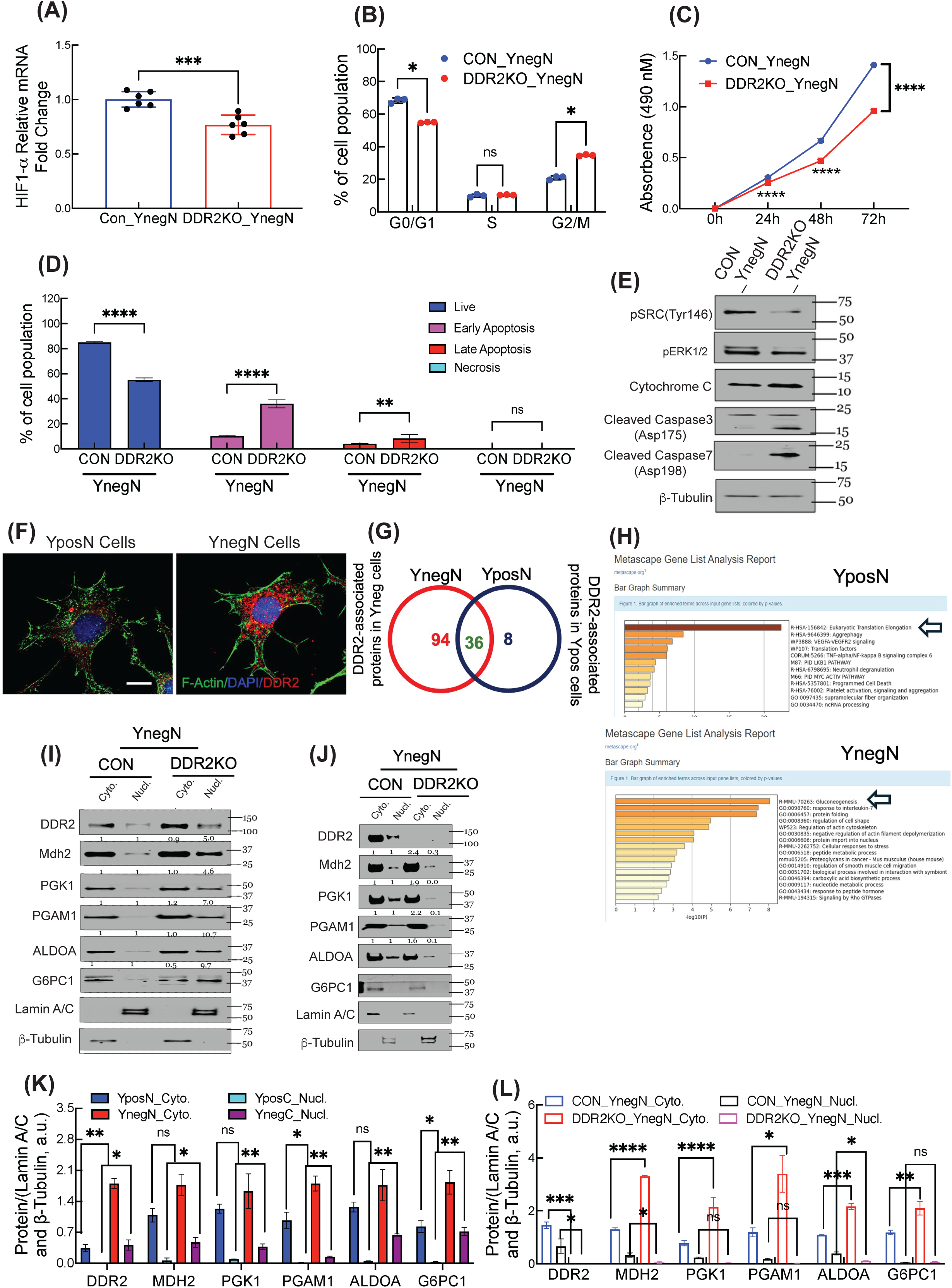
DDR2 interactome analysis. (A) HIF-1a gene expression analysis by qPCR. (B) Cell cycle analysis of CON_YnegN and DDR2KO_YNEGN cells; stage specific cell percentage quantitation. (C) MTS cell viability assay of CON_YnegN and DDR2KO_YNEGN cells, relative absorbance is plotted up to 72h. (D) Quantitation of cell percentages in specific apoptotic stage of for CON_YnegN and DDR2KO_YnegN cells from AnnexinV/PI apoptosis assay. (E) Immunoblot analysis of apoptotic pathway specific protein and phosphoproteins. (F) Immunofluorescence images of YposN and YnegN cells with DDR2. (G) Ven diagram showing the differential DDR2 interactomes in YposN and YnegN cells. (H) Metascape gene list analysis report for processes and pathways that includes the DDR2 interactomes found in YposN and YnegN cells. (I and K) Immunoblot analysis of spatial distribution of DDR and metabolic enzymes in cytoplasm and nucleus of YposN and YnegN cells, and quantitation (K). (J and L) Immunoblot analysis of spatial distribution of DDR2 and metabolic enzymes in cytoplasm and nucleus of CON_YnegN and DDR2KO_YnegN cells, and quantitation (L). Scale bars: 20 mM (F). Significance was calculated using either pairwise comparison (t-test) with welch correction (A, C) or with 2-way ANOVA with multiple comparisons (D, K, L): *p < 0.05, **p < 0.01, ***p < 0.001, and ****p < 0.0001.

### DDR2 complexes with gluconeogenic enzymes in LOY cells

DDR2 protein expression was increased in LOY cells, including prominent abundance in the nucleus (**Fig. 3B and S3A,B**), which is also noted by immunofluorescence (**Fig. 4F**). Next, to investigate DDR2-interacting proteins, we performed DDR2-immunoprecipitation (bait) in YposN and YnegN cells and analyzed the DDR2 interacting proteins by LC-MS. We found that DDR2 had ∼3-fold higher protein association with YnegN (130 proteins) compared to YposN (44 proteins) with 36 proteins common between both cell lines (**Fig. 4G and S4G**). DDR interactomes in both cell types by LC-MS were subjected to a system-level integrated analysis (**Fig. 4H**) to infer the enriched biological pathways and process using Metscape analysis (*43*). YposN cells had enrichment in eukaryotic translation elongation while YnegN cells had gluconeogenesis as the top-most enriched pathway with 5 gluconeogenic enzymes, Mdh2, Pgam1, Aloda1, Pgk1 and G6pc1 interacting with DDR2. YnegN cells also had enrichment of nuclear protein export, cytoskeleton remodeling, and protein folding (**Fig. 4H**). To explore this DDR2-enzymes association further, we first analyzed their respective localization in YnegN cells. Surprisingly, given that gluconeogenesis predominantly occurs in the cytoplasm (*44*), we also found these enzymes in the nuclear fraction (**Fig. 4I,K**). To evaluate if their interaction with DDR2 caused their nuclear presence, we determined their relative abundance in DDR2KO_YnegN cell (**Fig. 4J,L**) and found that compared to CON_YnegN cell, DDR2 deletion reduced their nuclear localization, suggesting their nuclear localization is DDR2 dependent.

## Discussion

Investigations into the clinical and biological relevance of tumor cell loss of the Y chromosome (LOY) have recently gained attention (*2, 4, 33*). Here, we examined whether changes in cellular metabolism could explain enhanced proliferation, stemness, and immune evasion associated with LOY. Normally, gluconeogenesis maintains glucose during fasting, while glycolysis operates under anaerobic conditions. Cancer cells suppress gluconeogenic enzymes like PEPCK and G6Pase, promoting glycolysis even in oxygen-rich settings (*45*), which supports fast ATP generation and biosynthesis (*12*). Since glycolysis and gluconeogenesis are reciprocally regulated, active gluconeogenesis would reverse glycolytic intermediates back to glucose, wasting energy. Proliferative cancer cells restrict gluconeogenesis to continue the aerobic glycolysis to meet their energy needs and biomass synthesis from glucose and glycolytic intermediates for growth and survival (*46*). The simultaneous operation of gluconeogenesis and glycolysis is considered a “futile cycle”, because it would lead to the consumption of ATP without any net production of glucose or other useful biomass for growth and proliferation, as the pathways would be effectively canceling each other out. In LOY cells, we speculate that DDR2 perhaps ensures that glycolysis and gluconeogenesis are not active simultaneously by translocating these enzymes to nucleus, so that cell maintains continuous aerobic glycolytic flux for sustained growth and proliferation (*47*). In DDR2KO_YnegN cells, when key enzymes failed to relocate in the nucleus, gluconeogenesis was active, evident in the increase of intracellular glucose level, and reduced lactate in culture media. Also, there was a reduction in non-glycolytic metabolites (amino acids, polyamines, fatty acids, lactate etc.) which are the substates for gluconeogenesis.

Cancer and immune cells compete for nutrients like glucose and amino acids, affecting tumor progression and immune function (*48*). Cancer cells prioritize glycolysis, while resting T cells rely on OXPHOS and fatty acid oxidation. Upon activation, T cells shift to glycolysis, increasing glucose demands. In tumors, cancer cells often outcompete T cells for glucose, impairing T cell function and reducing therapy efficacy. In melanoma, this metabolic competition leads to T cell starvation. Notably, transient glucose restriction can enhance CD8+ T cell activity, indicating a complex metabolism-immune relationship (*49*). Despite increased proliferation in YnegN cells, earlier, we found the viability remained unchanged in 2D cultures (*2*). This suggests highly proliferative LOY cells may also be dying at a higher rate, maintaining net population size. In culture, more floating cells were observed in YnegN versus YposN, indicating higher cell death. Proliferation appears localized to stem cell pools replenishing dying cells. We found DDR2 inhibition reduced aerobic glycolysis and lactate production in LOY cells. DDR2, known to activate HIF-1α (*50*), upregulates glycolytic enzymes and glucose uptake (*40*). While HIF-1α expression was not different between LOY and CON cells, DDR2 inhibition reduced HIF-1α, affecting glycolysis.

Emerging studies link DDR2 to metabolic reprogramming through interactions with metabolic enzymes and their subcellular translocation (*26, 51, 52*). We identified DDR2 interactions with glycolytic enzymes and their nuclear translocation, suppressing gluconeogenesis. Subcellular localization of metabolic enzymes supports tumorigenesis. For instance, mitochondrial translocation of Hexokinase 2 (HK2) helps evade apoptosis (*27, 53*). Pyruvate Kinase M2 (PKM2) moves to the nucleus to coactivate HIF-1α and c-Myc, promoting glycolysis (*54*). Nuclear translocation of PFKFB3 enhances glycolytic enzyme expression (*55*), while LDHA promotes lactate production and acidification (*56*). DDR2 also facilitates nuclear translocation of gluconeogenic enzymes like ALDOA1, MDH2, PGAM1, and PGK1. These enzymes, beyond their metabolic roles, can function as transcription factors or regulators, influencing gene expression (*57*). Whether DDR2-driven nuclear translocation solely suppresses gluconeogenesis or also modulates transcription relevant to LOY proliferation and immune evasion remains to be explored. Though dissecting these mechanisms is beyond our study, enrichment analysis of DDR2 interactome via LC-MS in LOY cells identified “protein import to nucleus” as an enriched pathway warranting further investigation.

In conclusion, our study maps the metabolic consequences of LOY in bladder cancer. Multiomics, imaging, and biochemical data suggest that LOY-associated aggressiveness and immune evasion are linked to metabolic reprogramming. DDR2 appears central to these shifts, and this suggests its inhibition presents a promising therapeutic avenue in LOY tumors.

## Materials and Methods

### Cell culture

The original parental MB49 cell line was a gift from M. Burdick. The MB49-YposN (cells containing Y chromosome genes) and YnegN cell lines (cells naturally lacking Y chromosome genes, hence calling YnegN, also known LOY; loss of Y cells) were cultured in DMEM supplemented with 10% heat-inactivated fetal bovine serum (FBS) and 0.1% sodium pyruvate. Single-cell clone isolation for YposN and YnegN cells was performed using serial dilution. Further, we have used CRISPR–Cas9-mediated Y chromosome knockout MB49 YposN cells; generated as described previously (*2*) (named as YposC cells; with intact Y chromosome and YnegC cells; without Y chromosome). Both cells were cultured in DMEM+10%FBS+10%Sodium Pyruvate in 37 °C, 5% CO_2_ environment. To generate DDR2 Knockout (DDR2KO_YnegN) cells through the CRISPR-Cas9 approach, 3 knockout plasmids targeting DDR2 were obtained. MB49 YnegN were transfected with three micrograms of plasmid DNA mixed with 150 µL of a diluted reagent, following the manufacturer’s protocol. Twenty-four hours after transfection, the transfection medium was removed and replaced with complete DMEM media. The cells were then maintained for an additional 72-hour post-transfection. After this incubation period, single cells exhibiting GFP (Green Fluorescent Protein) expression were sorted using flow cytometry in 96-well plates. The single-cell clones were pooled based on their expression levels, and immunoblotting was conducted to verify the results. Non-targeting CRISPR/Cas9 plasmids were used as a control.

### Immunoblotting of Nuclear Extracts and Total cell lysates

According to the manufacturer’s instructions (CHEMICON’s Nuclear Extraction Kit and NE-PER™ Nuclear and Cytoplasmic Extraction Reagents,) pure cytoplasmic fractions were extracted from the cellular lysates with cytoplasmic lysis buffer with protease, phosphatase inhibitors, DTT, and RNAase. Adherent cells were scraped and washed with PBS, and PBS was discarded by centrifugation at 250 g for 5 minutes at 4°C. After 15 minutes of incubation with 5-cell pellet volumes of ice-cold cytoplasmic lysis buffer, the supernatant was separated by centrifugation at 250 g for 5 minutes at 4°C. The supernatant was discarded, and the remaining pellet was resuspended in two ice-cold cytoplasmic lysis buffer volumes. Using a 27-gauge needle, the cell suspension was generated and centrifuged at 8000 g for 20 minutes at 4°C. The supernatant contains the cytosolic fraction of the cell lysates that were snap-freezed and stored at −80°C. The remaining pellet was resuspended in 2/3 of ice-cold nuclear extraction buffer protease, phosphatase inhibitors, DTT, and RNAase. Then, using a 27-gauge needle, nuclei were disrupted, and the nuclear suspension was gently agitated at 4°C for 30-60 minutes. The nuclear fraction was separated by centrifuging at 16000 g for 5 minutes at 4°C. The supernatant contains a nuclear fraction, and it is stored at −80°C. The total or fraction lysates were separated by 10-15% SDS-PAGE and transferred onto the PVDF membrane by BioRad semi-dry transfer system for 10 minutes. After the transfer, the membranes were blocked with 5% nonfat dry milk (NFDM) in TBST (20 mM Tris, pH 7.6, 100 mM NaCl, 0.1% Tween-20) for 1 h at room temperature. Then, the membranes were rinsed with TBST and incubated with primary antibodies in 5% NFDM or BSA overnight at 4°C. Afterward, the membranes were washed thrice with TBST and incubated with secondary antibodies (1:5000 or 1:10000), conjugated with horseradish peroxidase (HRP) in 1% BSA or NFDM in TBST for 1 h at room temperature. Following this, the membranes were washed thrice with TBST for 10 minutes, and signals were detected by Pierce™ ECL Western Blotting Substrate (ThermoFisher Scientific: Cat. No.32106). Antibodies were used included following: DDR2 (Cell Signaling Technology, 12133S), DDR2 (E5S1S; Rabbit mAb #71991), DDR2 Polyclonal Antibody (PA5-27752; Thermo), Lamin A/C (Cell Signaling Technology, 4777S), Lamin B1 (Cell Signaling technology), GAPDH (14C10; Rabbit mAb #2118; Cell Signaling technology), β-Tubulin (Cell Signaling Technology, 2128S), β-Actin (Cell Signaling Technology, 4970S), Human Phospho-DDR1/DDR2 (R&D Systems, MAB25382-SP), Mdh2 (Abcam, ab96193), PGK1 (Proteintech, 17811-1-AP), PGAM1 (Proteintech, 16126-1-AP), ALDOA (Proteintech, 11217-1-AP), G6PC (Proteintech, 66860-1-Ig), Src (Cell Signaling Technology, 2109S), p-Src(Tyr416) (Cell Signaling Technology, 6943S), Erk1/2 (Cell Signaling Technology, 4695S), pErk1/2 (Thr202/Tyr204) (Cell Signaling Technology, 4370S), Cytochrome c (Cell Signaling Technology, 11940S), Cleaved Caspase-3 (Asp175) (Cell Signaling Technology, 9661S), Cleaved Caspase-7 (Asp198) (Cell Signaling Technology, 8438S), Goat anti-Mouse IgG (H+L) secondary antibody HRP (Fisher Scientific, PI32430), Goat anti-Rabbit IgG (H+L) secondary antibody HRP (Fisher Scientific, PI32460).

### Immunofluorescence

YposN (or YposC) and YnegN (or YnegC) cells were seeded onto glass coverslips coated with 1% poly-L-lysine and allowed to attach and divide overnight. The following day, the cells were fixed with 4% paraformaldehyde and permeabilized using Triton X-100. The primary antibody for DDR2 (Proteintech, 67126-1-Ig) was incubated overnight at 4°C and then washed three times with PBS for 10 minutes each time. DDR2 was visualized using an Alexa Fluor 555-conjugated goat anti-mouse secondary antibody (Abcam, ab150114) at room temperature for 1 hour. Filamentous actin (F-actin) was stained with FITC-Phalloidin (Life Technologies, A12379) for 30 minutes. After two additional washes with PBS, DRAQ5 (Thermo Scientific, 62251) was employed to stain nuclei in the dark for 10 minutes. The coverslips were then air-dried and mounted with ProLong Gold Antifade (ThermoFisher Scientific, P10144). Finally, the subcellular distribution of DDR2 was analyzed using a Leica Stellaris 8-STED Super-resolution Confocal Microscope, and the images were evaluated with Leica Application X software.

### UHPLC-MS based untargeted metabolomics: experiment and data analysis

All cell lines were cultured for multiple passages with supplemented 10% FBS and 1% penicillin streptomycin. Then the lysates were made by RIPA lysis buffer with protease, phosphatase II and phosphatase III inhibitors. After the protein quantification 1mg of protein was used for DDR2 immunoprecipitation. After this incubation period, the supernatant was carefully removed, and the cells were washed three times with cold phosphate-buffered saline (PBS). Cell lysis was performed using 100μL of 8M Urea in 50mM Tris-HCl (pH 8.0). The lysates were then transferred to pre-chilled 1.5mL microcentrifuge tubes and centrifuged at 15,000 RCF for 10 minutes to pellet debris. The resulting supernatant was collected in a fresh, pre-chilled tube and stored frozen. The protein aliquots were digested and cleaned using the S-Trap system (Protifi). The elution from the Strap columns were dried and resuspended at a concentration of 1 µg/µl for injection onto mass spectrometry (*31*) system. To extract metabolites, cold 5:3:2 MeOH:ACN:H2O (v/v/v) solution was added to cells in a final concentration of 2 million cells/mL. Samples were then vortexed vigorously for 30 minutes at 4°C, then centrifuged for 10 minutes at 18,213 rcf. Using 10 µL injection volumes, the supernatants were analyzed by ultra-high performance liquid chromatography coupled to mass spectrometry (UHPLC-MS — Vanquish UHPLC Autosampler and Exploris 120 Orbitrap mass spectrometer, ThermoFisher). Metabolites were resolved across a 1.7 um, 2.1 x 150 mm Kinetex C18 column using a 5-minute gradient previously described (*58*). Following data acquisition, .raw files were converted to .mzXML using RawConverter. Metabolites were then assigned based on intact mass, 13C isotope pattern and retention times in conjunction with the KEGG database and an in-house standard library. Peaks were integrated using Maven (Princeton University). Quality control was assessed as using technical replicates run at beginning, end, and middle of each sequence as previously described (*59*). Raw peak data was normalized by median and auto-scaled then was used to generate figures in MetaboAnalyst (*60*). Intensity values were log2 transformed and p-values were calculated using two-tailed t-tests assuming equal variance. Adjusted p-values were calculated by Benjamini-Hochberg correction. Log2FoldChange values were calculated based on mean of the log2 transformed intensity values. Volcano plots were generated based on log10(padj) and log2FoldChange values.

### Co-Immunoprecipitation of DDR2

The cell lysates were prepared by lysing the cells in RIPA lysis buffer supplemented with protease inhibitor cocktail (Sigma-Aldrich, Cat. No. P8340), Phosphatase inhibitor cocktail 2 (Sigma-Aldrich, Cat. No. P5726) and phosphatase inhibitor cocktail 3 (Sigma-Aldrich, Cat. No. P0044) for 30 min at 4oC. After the Bradford protein assay (Bio-Rad, CA. No. 5000006), 500 to 3000µg of total protein lysate was used in immunoprecipitation (IP) reactions. IP was carried out using Protein G Sepharose beads following the manufacturer’s protocol (GE HealthCare). The protein lysates were incubated with mouse IgG monoclonal *DDR2* antibody (Proteintech, 67126-1-Ig) at 4°C overnight with gentle mixing end over end shaker. The next day, the slurry of sepharose beads was washed thrice with the lysis buffer by centrifuging at 2000*g* for 2 min at 4°C. After washing three times with a lysis buffer, the mixture was incubated with the protein lysates and DDR2 antibody at 4°C for 4 hours by gentle mixing end over end shaker. Next, the precipitated protein with the bead was washed three times and analyzed using immunoblotting. Then the immunoprecipitated DDR2 was blotted with rabbit Polyclonal MDH2 antibody (1:5000, Abcam, ab96193), rabbit Polyclonal Pgk1 antibody (1:5000, Proteintech, 17811-1-AP), rabbit Polyclonal Pgam1 antibody (1:5000, Proteintech, 16126-1-AP), rabbit Polyclonal Aldoa1 antibody (1:5000, Proteintech, 11217-1-AP).

### LC/MS analysis of DDR2-coIP: experiment and data analysis

After performing DDR2 immunoprecipitation, the agarose beads were incubated in 8 M urea with 50 mM Tris-HCl (pH 8.0). The lysates were then transferred to pre-chilled 1.5 mL microcentrifuge tubes and centrifuged at 15,000 RCF for 10 minutes to remove debris. The resulting supernatant was carefully collected into a fresh pre-chilled tube and stored at freezing temperatures. Next, the samples were treated with 10 mM DTT for reduction, followed by alkylation with 100 mM IAA. The pH of the diluted protein lysates was adjusted to a range of 7 to 8, and enzymatic digestion was carried out using sequencing-grade Trypsin/Lys-C (Promega) in the presence of 10% acetonitrile for 16 hours at 37°C. After digestion, the samples were acidified by adding formic acid to a final concentration of 1%, and any residual acetonitrile was removed through speed-vac dehydration. The peptides were then desalted using C18 tips (Nest Group) and thoroughly dried. Before mass spectrometry analysis, the peptides were reconstituted in a 0.1% formic acid solution at a concentration of 0.5 μg/μL, with synthetic iRT reference peptides (Biognosys) added at a 1:40 ratio. For data acquisition, Following established protocols, peptides were analyzed using liquid chromatography coupled with mass spectrometry in data-independent acquisition (DIA) mode. Briefly, 4 μL of the digested sample was directly injected onto a 200 cm micro-pillar array column (uPAC, Pharmafluidics) and separated over a 120-minute reversed-phase gradient at a flow rate of 1200 nL/min and a temperature of 60°C. The gradient consisted of aqueous 0.1% formic acid (solvent A) and 0.1% formic acid in acetonitrile (solvent B), applied as follows: 2% B from 0 to 5 minutes, increased to 4% B at 5.2 minutes, then linearly ramped to 28% B by 95 minutes, and further increased to 46% B at 120 minutes. After each analytical run, the column was flushed at 1200 nL/min and 60°C by injecting 50% methanol at 95% B for 25 minutes, followed by a 10-minute ramp down to 2% B and a 5-minute equilibration at 2% B. The eluting peptides were electrosprayed through a 30 μm bore stainless steel emitter (EvoSep) and analyzed on an Orbitrap Lumos mass spectrometer using DIA within the 400–1000 m/z range.

### Total-RNAseq: experiment and data analysis

Total RNA was extracted using the RNeasy Plus Kit (Qiagen), following the manufacturer’s instructions. Y-CRISPR control and DDR2 knockout cells were lysed and homogenized using a QIAshredder column (Qiagen), with three distinct biological replicates included. To eliminate any remaining genomic DNA, gDNA elimination columns (Qiagen) were used. The quality and quantity of the extracted RNA were assessed using a NanoDrop spectrophotometer, agarose gel electrophoresis, and an Agilent 2100 Bioanalyzer.

For processing the sequencing data, Illumina TruSeq adapter sequences, as well as polyA and polyT sequences, were trimmed using Cutadapt v2.3. The processed reads were then aligned to the mouse genome (mm10, version 38) using STAR aligner v2.7.0d, in accordance with the recommendations of the ENCODE long RNA-seq pipeline (https://github.com/ENCODE-DCC/long-rna-seq-pipeline). Gene expression levels were quantified with RSEM v1.3.1 using Ensembl gene annotations (version 84) for both alignment and quantification. The quality of the RNA-seq data, which includes metrics on sequencing, alignment, and quantification, was evaluated using FastQC v0.11.5 and MultiQC v1.8. Principal component analysis (PCA) and pairwise Pearson correlation analysis were performed to assess the concordance of biological replicates. Lowly expressed genes were filtered out based on the criterion that estimated counts from RSEM must be at least five times the number of samples. Differential expression analysis was conducted using the DESeq2 v1.22.2 package from R Bioconductor, employing a generalized linear model with a negative binomial distribution. The Benjamini–Hochberg method was applied for multiple testing correction.

### Colony Forming Assay

CRISPR control (CON_YnegN) and DDR2 knockout cells (DDR2KO_YnegN, 100 per well) were seeded into a 6-well plate containing DMEM medium supplemented with 10% FBS and 1% penicillin-streptomycin. After incubating for 7 days, the cells were washed with PBS and fixed using 100% methanol for 20 minutes. Once the methanol was removed, the cells were rinsed with purified water. A crystal violet staining solution was then added in a sufficient volume and incubated for 5 minutes at room temperature. The cells were subsequently washed with water until the excess dye was removed. Images were captured using a bright-field microscope, and colony quantification was performed using ImageJ software. This experiment was repeated three times with cells from different passages.

### Apoptosis Assay

CON_YnegN and DDR2KO_YnegN cells were seeded in a 10 cm plate and incubated for 24 hours. After incubation, the cells were collected by centrifugation, washed with PBS, and stained using the FITC Annexin V Apoptosis Detection Kit (BioLegend, San Diego, CA, USA) according to the manufacturer’s instructions. Flow cytometry analysis was performed using the Fortessa, Sony SA3800 Cell Analyzer (BD Biosciences, Vaud, Switzerland), and the data were processed with FlowJo™ v10.6.1 (BD Life Sciences). The experiments were conducted in triplicate, a practice that was consistently followed to ensure the reliability of the results. Annexin V-positive and 7-AAD-negative cells were classified as apoptotic, while Annexin V-positive and 7-AAD-positive cells were considered late necrotic. Flow cytometry analysis was performed using the BD LSR Fortessa™ Cell Analyzer (BD Biosciences, Vaud, Switzerland), and data were processed with FlowJo™ v10.6.1 (BD Life Sciences). Experiments were conducted in triplicate, further reinforcing the reliability of the results. Annexin V-positive, 7-AAD-negative cells were classified as apoptotic, while Annexin V-positive, 7-AAD-positive cells were considered late necrotic.

### Cell proliferation assay

Cell proliferation was evaluated using the CellTiter 96 AQueous One Solution Assay (Promega), following the manufacturer’s instructions. Approximately 5,000 cells were seeded in each well of a 96-well plate for appropriate time, and 20 µL of the reagent/100 µL culture medium was added to every well. The plate was incubated for one hour in a humidified atmosphere containing 5% CO. After incubation, absorbance was measured at 490 nm every 24 hours using a SpectraMax I3 microplate reader (Molecular Devices, Sunnyvale, CA). Measurements were taken from six replicate wells for each time point.

### Cell cycle analysis

Cell cycle analysis was performed using Propidium Iodide (Thermo Scientific; P1304MP) as per manufacturer’s recommended protocol. Briefly, cells were grown in DMEM media in 6-well plates for appropriate time and conditions. Afterwards, cells were trypsinized and floating cells were also collected for the analysis and fixed in 70% chilled ethanol. Cells were washed twice with PBS, stained with Propidium Iodide (100 µg/mL) along with RNAase-A (100 µg/mL), and acquisition was performed through a BD Biosciences Symphony A5 Cell Analyzer. Two color compensation was performed for Propidium Iodide and GFP.

### Glucose and Lactate assay

Glucose and L-lactate abundance in the culture media, or within the clarified cell lysate, were analysed using RayBiotech colorimetric assay kit (MA-GLU-1 and MA-LAC-1 respectively), and measured as recommended in manufacturer’s protocol. Approximately 1×10^5^ cells were seeded in each well of a 6-well plate for 72h before the culture media or cell lysate obtained for analysis.

### TCGA Analysis

Total-RNA seq data for bladder cancer in the Pan-Cancer Atlas was downloaded from the GDC portal using project id (TCGA-BLCA). Normalized counts were calculated using the DESeq2 package and patients without survival data were excluded from the analysis. Normalized TCGA bulk RNA Seq data counts were log-transformed (log2(1+exp)) and median centered. z-scores based on the expression of all the Y genes were calculated for each patient similar to the “Z-score activation metric” described in Levine *et al. 75.* Next, surv_cutpoint function from survminer package was used to stratify patients into high and low expression groups. survival and survminer packages were used to perform survival analysis and univariate cox regression analysis was used to determine statistical significance and hazard ratio. The patients from TCGA were thus classified into Yhigh and Ylow groups and box plots of DDR2 expression were generated.

### Pan-cancer scRNA-seq Data Collection

For pan-cancer single-cell RNA sequencing (scRNA-seq) data, we utilized data compiled in our previous work (*33*). This dataset comprises transcriptomic profiles from 346 samples across 251 individuals, collected from 20 publicly available scRNA-seq studies. Tumor samples were selected from this dataset for downstream analysis. To integrate and correct for batch effects in the scRNA-seq data, we employed the scVI Python package (scvi-tools, version 1.0.4) (*61*). Cell type annotation and quality control procedures followed the protocols established in our previous study. For functional signature calculation for scRNA-seq Data, we computed functional gene set scores using the scanpy.tl.score_genes function from the Scanpy Python package (version 1.9.5). This function assigns a score to each cell based on the expression of genes in a specified list. The glycolysis gene set was downloaded from the GSVA database (https://www.gsea-msigdb.org/).

### Real-Time qPCR

Total RNA was extracted from YposN (or YposC) and YnegN (or YnegC) using RNeasy Mini Kit per the manufacturer’s protocol. cDNA was synthesized from the isolated RNA using a SuperScript III first-stand synthesis kit. Afterward, real-time PCR was performed using 2X PowerUp SYBR Green Master Mix. In the PCR reaction, 2 µL of cDNA and 2 µL of forward and reverse primers were used. Quantification was performed using the ΔΔCt method and β-actin and Gapdh primers as the control. The values from Ypos cells were normalize to 1. Similarly, RNA was extracted from CON_YnegN and DDR2KO_YnegN cells using RNeasy plus Mini Kit and cDNA was synthetized using Maxima H Minus cDNA Synthesis Master Mix (Thermo Scientific) as per manufracturer recommended protocol. Subsequently, qPCR was performed with powerUp SYBR Green Master Mix (Applied Biosystems) on Quant Studio 6 Flex Real-Time PCR system (Applied Biosystems). Changes in mRNA were quantified using ΔΔCt method using GAPDH as internal control. qPCR primer sequences are provided in supplementary table S3.

### Flow cytometry analysis for quantification of CD33, CD44, and Ki67 levels in cells

YposC and YnegC cells were grown in DMEM+10%FBS+10%Sodium Pyruvate in 37 °C, 5% CO_2_. Cells were trypsinized and passed through filter capped flow cytometry tubes to make single-cell suspension. Cells were washed and surface staining was conducted with CD44 antibodies (CD44 Monoclonal Antibody (IM7), FITC, eBioscience, Cat. 11-0441-8211-0441-82; PE/Cyanine7 anti-mouse/human CD44 Antibody Biolegend, Cat. 103029) and APC anti-mouse CD133 (Biolegend, Cat. 141208) antibodies alone and in combination. Iso-FITC, Iso-PE/Cyanine7 and unstained cells were used as controls, for combination staining a two-color compensation was performed for acquisition through a BD Biosciences Symphony A5 Cell Analyzer. For single staining, CD44-positive cells and CD133-positive were gated to measure percentage cell population of CD44+ and CD133+ cells. For double staining group, live cells were gated, CD44-positive cells, CD133+ cells, and CD44+ CD133+ (double-positive) cells were calculated, and percentage mean fluorescence intensity (MFI) was determined for each population. For Ki-67 analysis, cells were trypsinized and passed through filter capped flow cytometry tubes, fixed in 70% chilled ethanol and stained with BD Horizon™ BV786 Mouse Anti-Ki-67 antibody, unstained cells were used as control and acquisition was performed through a BD Biosciences Symphony A5 Cell Analyzer.

### Cell Proliferation analysis

For Ki67 based analysis, post-fixation (4% PFA) and permeabilization, cells were incubated with anti-Ki67 primary antibody (SolA15, ThermoFisher; 1:500 dilution) for 2h, and then incubated with secondary antibody and DAPI (10 mg/ml; 1:2000) for 30 minutes, cells were then mounted with Prolong-Gold and imaged for Ki67 positive nucleus and total number of nucleus in each segmented images. %-of Ki67 positive nuclei were plotted for comparison among samples. For, Edu-proliferation assay, the Click-iT™ EdU Cell Proliferation Kit for Imaging, Alexa Fluor™ 594 dye (ThermoFisher, Cat# C10339) was used, cells were prepared as suggested by manufacturer. Similar to Ki67, the %-of Edu positive cells were counted for samples and plotted for comparison.

### Statistical Analysis

The data analysis was performed by employing the GraphPad Prism version 10.0 software. Unpaired t-test, and 2-way ANOVA analysis were performed (as mentioned in figure legends) to determine the statistical significance between the groups, as mentioned in figure legends. Three biological replicates were performed in all the experiments unless otherwise stated. All results are presented as means ± SEM, with P values and significance denoted in the figures [not significant (n.s.) > 0.05, *P < 0.05, **P < 0.01, ***P < 0.001, and ****P < 0.0001].

## Author contributions

Funding: DT; Conceptualization: DT, BRA, BS; Methodology: BS, PA, SKKM, SM, XC, HA, and BRA; Investigation: BS, PA, SKKM, SM, XC, AQ, YF, HA and BRA; Visualization: DT, BRA, SKKM, BS, PA, SM; Supervision: DT, BRA, JC; Writing original draft: BRA and DT; Writing review & editing: DT, BRA, BS, PA, SKKM, JC, XC.

## Data and materials availability

All data needed to evaluate the conclusions in the paper are present in the paper and/or the Supplementary Materials.

## Supplementary Materials

Supplementary file contains Figure S1 to S4.

**Figure S1.**
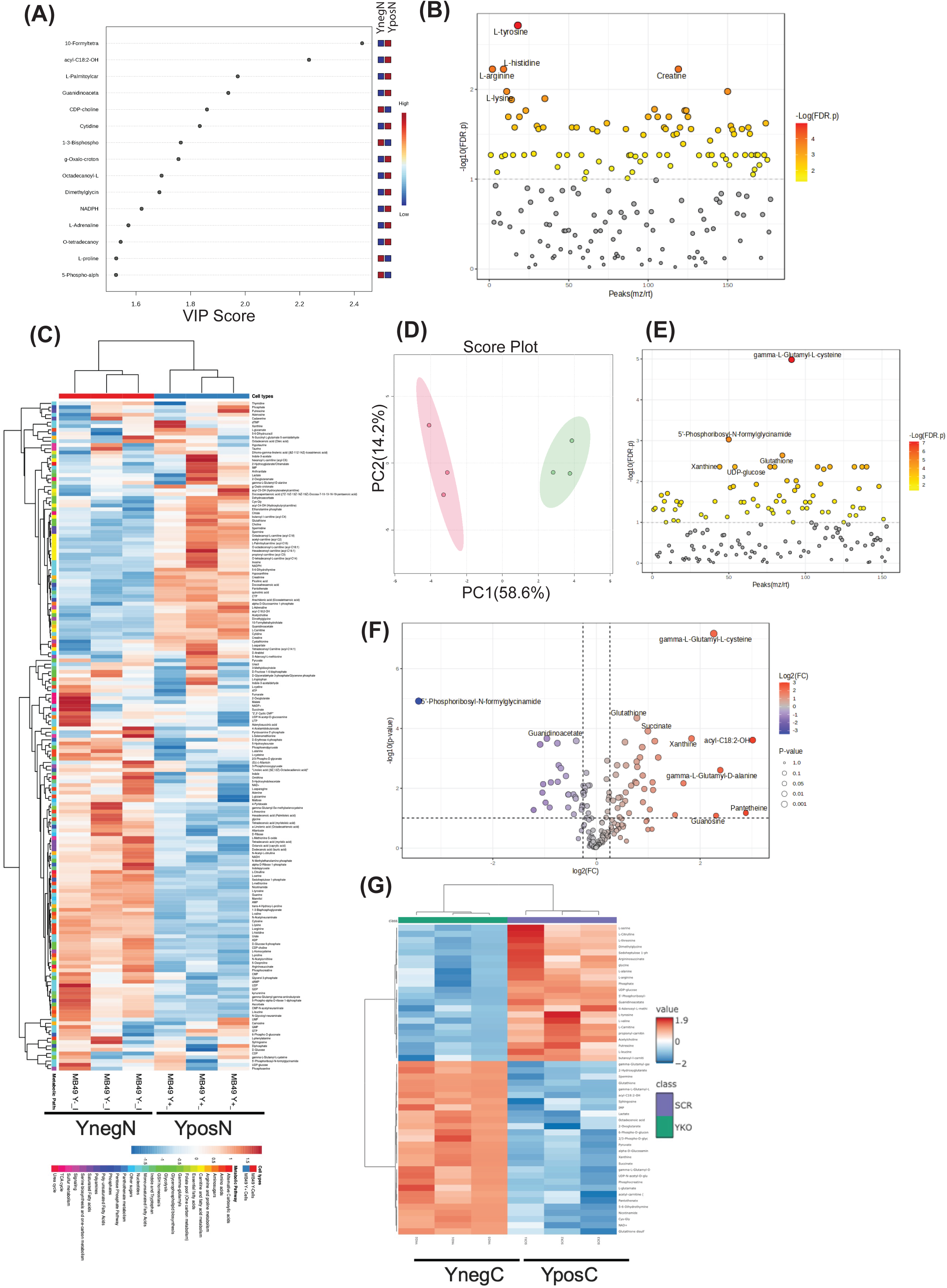

**Figure S2.**
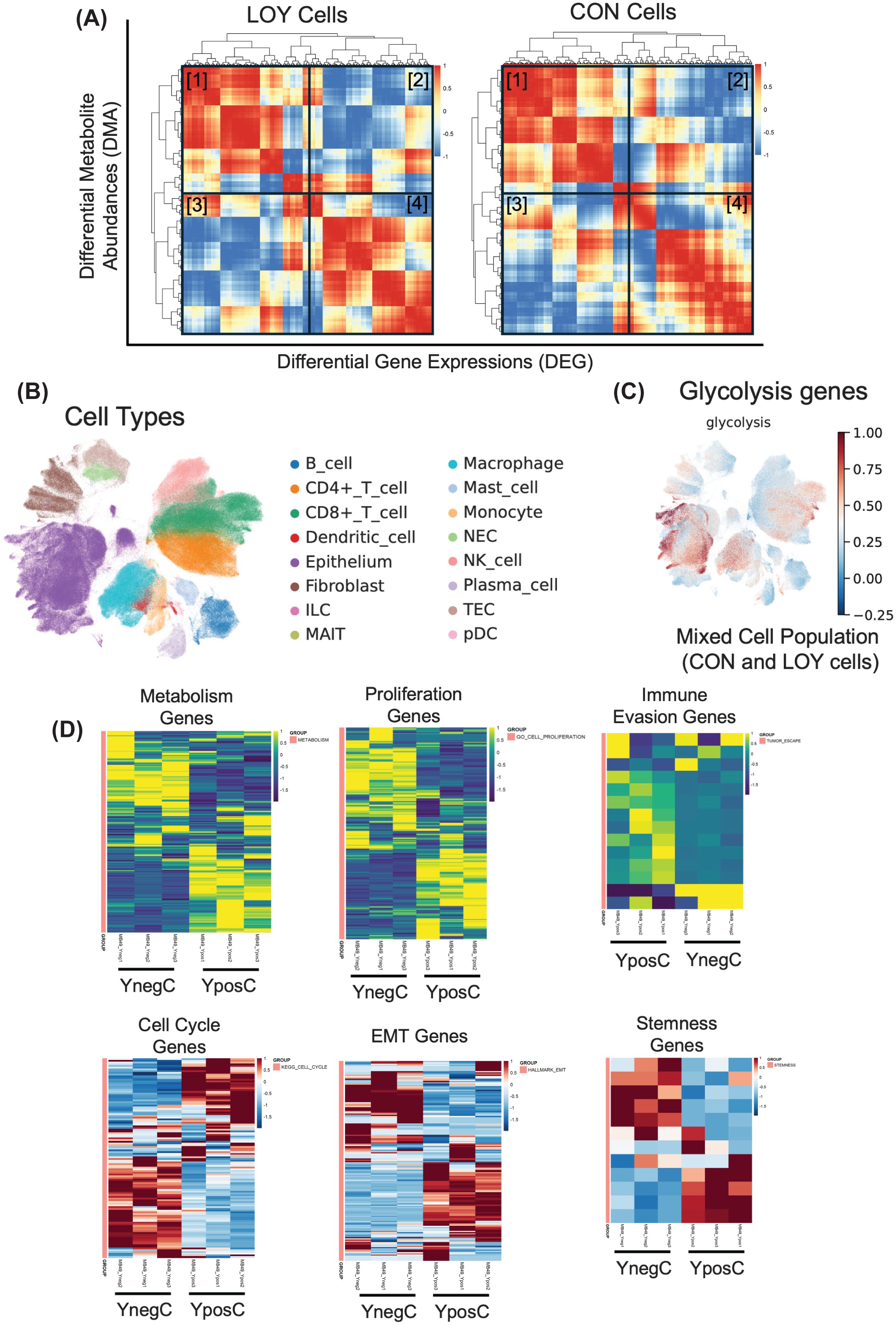

**Figure S3.**
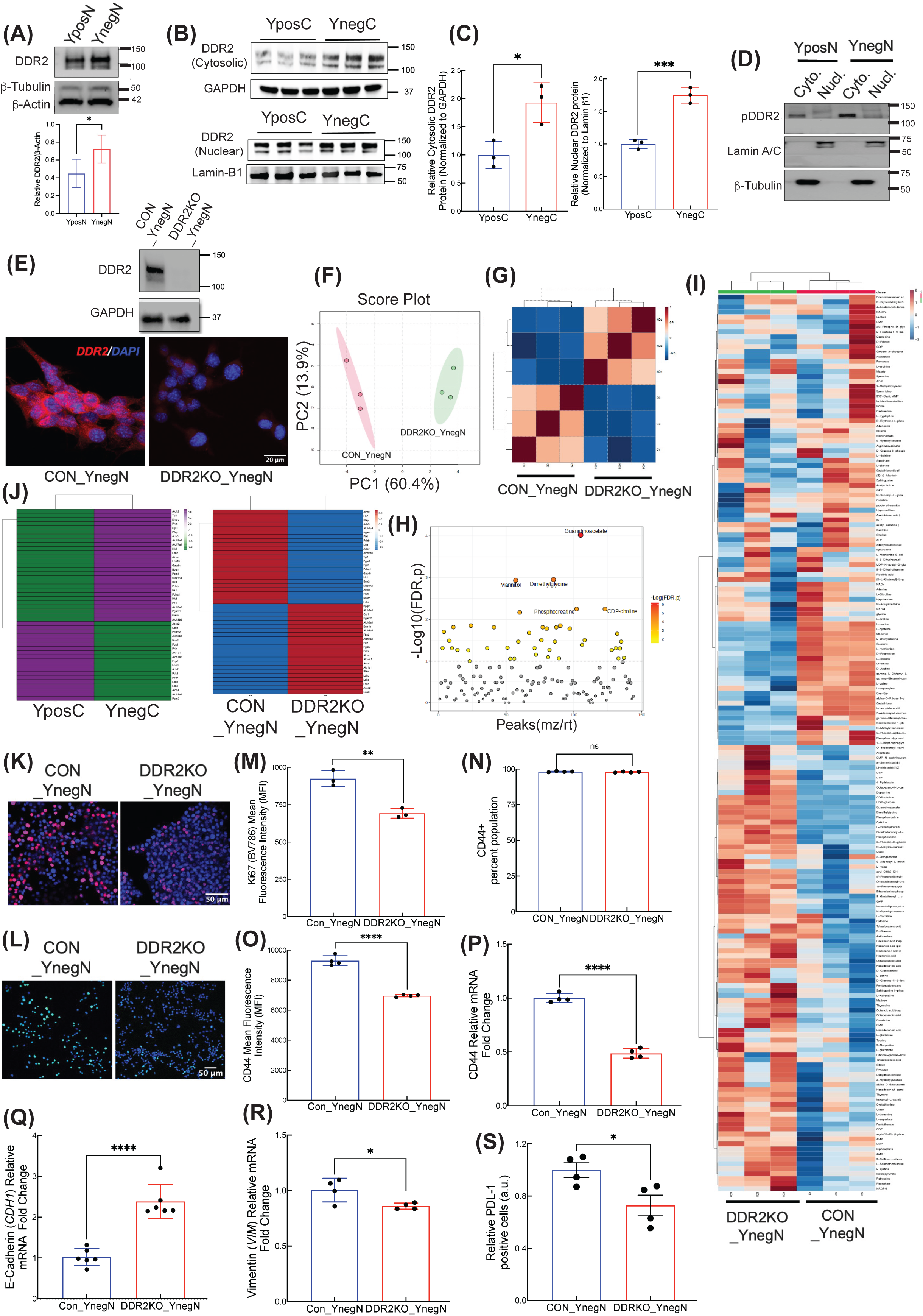

**Figure S4.**
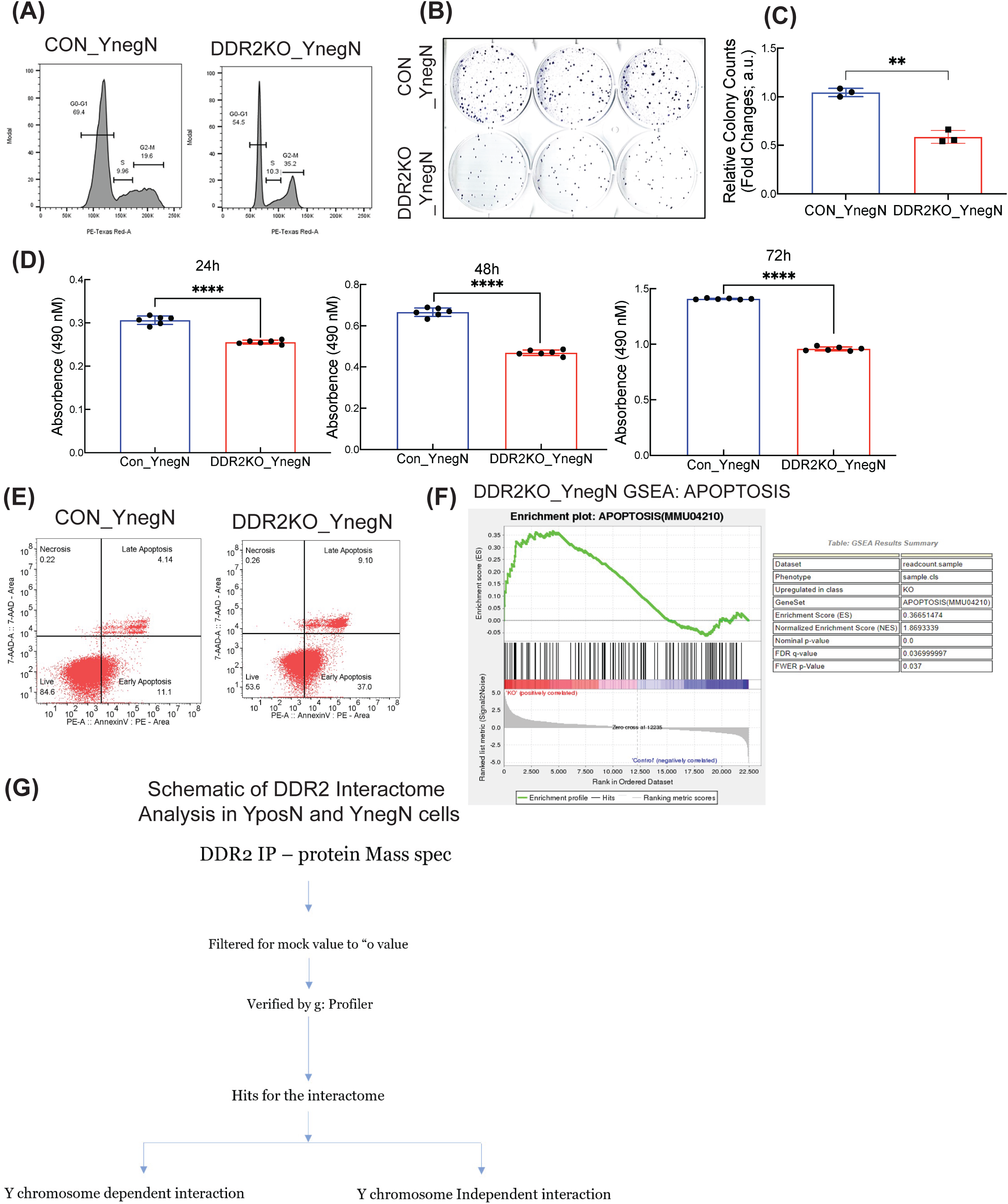

