## Supplementary text for "Loss of the Y chromosome in bladder cancer drives metabolic reprogramming"

Boopathi and Acharya *et al.*

**This file includes:**

Supplementary figures; Fig S1 to S4

Supplementary text: Figure legends corresponding to Fig S1 to Fig S4 Supplementary tables: Table S1: Chemical and Reagent list

Table S2: Assay kit list Table S3: qPCR Primer list

**Supplementary Text**

**SUPPLEMENTARY FIGURE LEGENDS**

**Figure S1:** (A) VIP plot for top10 metabolites causes the variations between YnegN and YposN cells. (B) Significant differential metabolites abundance (DMA) t-test plot, with p-value cut off at

0.1 and fold difference cut off at 1.2. (C) The Heatmap all DMA between Yneg vs Ypos belongs to multiple metabolism pathways. (D) PCA analysis showing variance between YposC and YnegC cell metabolome. (E) Significant differential metabolites abundance (DMA) in YposC vs. YnegC cells; t-test plot, with p-value cut off at 0.1 and fold difference cut off at 1.2. (F) Volcano plot analysis for significant metabolites between YposC and YnegC cells. with p-value cut off at 0.1 and fold difference cut off at 1.2. (G) Heatmap of 50 metabolites with highest fold differences in DMA of YnegC vs. YposC.

**Figure S2:** (A) Integrated differential metabolome and transcriptome correlation heatmaps for Ypos and Yneg cell for all DMA and all DEG. (B) Uniform Manifold Approximation and Projection (UMAP) visualization of 497,055 single cells from human tissues profiled by single- cell RNA sequencing. Left: cells are colored by manually curated cell type annotations across diverse lineages, including B cells, T cells (CD4⁺ and CD8⁺), dendritic cells, macrophages, epithelial cells, fibroblasts, and others (see legend for full list). (C) Same UMAP for all cells, regardless with or without Y chromosomes is colored by glycolysis gene set enrichment score (GSEA-derived or AddModuleScore), highlighting cell type–specific variation in glycolytic activity. Warmer colors indicate higher glycolytic activity. (D) The DEGs for 3 independent samples of Ypos and Yneg cells for specific genesets as indicated on top of the heatmap.

**Figure S3:** (A) Western blot analysis of total DDR2 in YposN and YnegN cells. (B and C) DDR2 protein abundance in cytoplasm and nuclear fraction of YposC and YnegC cells, and quantification

(C). (D) phospho-DDR2 (Tyr470) in cytoplasm and nuclear fraction in YposN and YnegN cells.

(E) Characterization of DDR2 Knock out in Yneg cells, immunoblot and immunofluorescence images. (F) PCA analysis showing variance between CON_YnegN and DDR2KO_YnegN cell metabolome. (G) Pearson Correlation heatmap of variance between CON_YNEGN and DDRKO_LOY cell. (H) Significant differential metabolites abundance (DMA) t-test plot, with p- value cut off at 0.1 and fold difference cut off at 1.2. (I) The Heatmap all DMA between

DDR2KO_YnegN vs CON_YnegN cells belongs to multiple metabolism pathways. (J) Normalized DEG prolife of KEGG glycolysis/gluconeogenesis genes for both YposC and YnegC cells and CON_YnegN and DDR2KO_YnegN cell types. (K and L) Immunofluorescence images of nuclei for Click_IT-Edu cell proliferation and Ki67 proliferation/division assay (L). (M) Mean- fluorescence intensity of Ki67 in DDR2KO_YnegN and CON_YnegN cells detected by flow cytometry. (N and O) Cell surface expression, either the % of cells (N) or mean fluorescence intensity (O) of cancer stem cell markers CD44 by flow cytometry and qPCR analysis. (P) CD44; cancer stem cell marker gene expression analysis by qPCR. (Q and R) EMT genes expression analysis by qPCR of E-cadherin (*CDH1*) and Vimentin (*VIM*) (R). (S) Surface expression of PD- L1 on CON_YnegN and DDR2KO_YnegN cells analyzed by flow cytometry. Scale bars: 20 mM (E), 50 mM (K, L). Significance was calculated using either pairwise comparison (t-test) with welch correction (C, M-S): *p < 0.05, **p < 0.01, ***p < 0.001, and ****p < 0.0001.

**Figure S4:** (A) Flow cytometry panels of Cell cycle analysis of CON_YnegN and DDR2KO_YnegN cells. (B and C) Clonogenic assay plate image (B) and colony quantitation (C) for CON_YnegN and DDR2KO_YnegN cells. (D) MTS cell viability assay quantitation plotted as cumulative data for each 24h until 72h. (E) Flow cytometry panel for AnnexinV/PI apoptosis assay for CON_YnegN and DDR2KO_YnegN cells. (F) GESA analysis of Apoptosis genes in DDR2KO_YnegN cells. (G) Schema of DDR2 protein-protein interaction analysis by LC/MS.

**Table S1: Chemicals and reagents**

| Item Name | Company | Catalog Number |
| --- | --- | --- |
| Corning™ DMEM (Dulbecco’s Modified Eagle’s Medium) | Fisher Scientific | MT10017CV |
| FBS QUALIFIED USA ORIGIN | Fisher Scientific | 26140079 |
| Gibco™ Sodium Pyruvate (100 mM) | Fisher Scientific | 11-360-070 |
| Halt™ Protease Inhibitor Cocktails | Fisher Scientific | PI78429 |
| Phosphatase inhibitor cocktail 2 | Sigma-Aldrich | P5726 |
| Phosphatase inhibitor cocktail 3 | Sigma-Aldrich | P0044 |
| Gibco DPBS without Calcium and Magnesium | Fisher Scientific | 14190144 |
| DTT | Sigma-Aldrich | SKU 10197777001 |
| Poly-L-lysine solution | Sigma Aldrich | P4832-50ML |
| Paraformaldehyde, 4% in PBS | Thermo Fisher Scientific | J61899.AK |
| Background Sniper | Thermo Fisher Scientific | 50-823-85 |
| Alexa Fluor™ 488 Phalloidin | Thermo Fisher Scientific | A12379 |
| DRAQ5™ Fluorescent Probe Solution | Thermo Fisher Scientific | 62254 |
| Molecular Probes™ ProLong™ Live Antifade Reagent, for live cell imaging | Fisher Scientific | P36975 |
| Pierce™ IP Lysis Buffer | Thermo Fisher Scientific | 62254 |
| Trypan Blue Solution, 0.4% | Fisher Scientific | 15250061 |
| Protein G Sepharose® 4 Fast Flow | Sigma Aldrich | GE17-0618-01 |
| Gibco Penicillin-Streptomycin (10,000 U/mL) | Fisher Scientific | 15140-122 |
| RIPA Lysis buffer | Thermo Fisher Scientific | 89900 |
| Propidium Iodide | Thermo Fisher Scientific | P1304MP |
| RNAse-A | Thermo Fisher Scientific | EN0531 |

**Table S2: Assay Kits**

| Item Name | Company | Catalog Number |
| --- | --- | --- |
| Nuclear Extraction Kit | EMD | 2900 |
| NE-PER™ Nuclear and Cytoplasmic Extraction Reagents | Thermo Scientific | 78833 |
| Pierce™ BCA Protein Assay Kit | Thermo Fisher Scientific | 23227 |

| Duolink® In Situ Wash Buffers, Fluorescence | Sigma Aldrich | DUO82049-4L |
| --- | --- | --- |
| Duolink® In Situ Detection Reagents Red | Sigma Aldrich | DUO92008-100RXN |
| Duolink® In Situ Red Starter Kit Mouse/Rabbit | Sigma Aldrich | DUO92101 |
| Control Double Nickase Plasmid | Santa-cruz | sc-437281 |
| DDR2 Double Nickase Plasmid (m) | Santa-cruz | sc-421985-NIC |
| RNeasy Plus Mini Kit (250) | Qiagen | 74136 |
| QIAshredder (250) | Qiagen | 665472 |
| FITC Annexin V Apoptosis Detection Kit with 7-AAD | Biolegend | 640922 |
| CellTiter 96® AQueous One Solution Cell Proliferation Assay (MTS) 5000 Assay Kit | Promega | G3581 |
| SuperScript™ III First-Strand Synthesis System | Thermo Fisher Scientific | 18080051 |
| PowerUp™ SYBR™ Green Master Mix for qPCR | Fisher scientific | A25742 |
| Maxima™ H Minus cDNA Synthesis Master Mix | Thermo Fisher Scientific | M1662 |

**Table S3: qPCR Primers**

| Gene | Forward Primer | Reverse primer |
| --- | --- | --- |
| Gapdh | AACAGCAACTCCCACTCTTC | CCTGTTGCTGTAGCCGTATT |
| Aldh1a1 | GGAATACCGTGGTTGTCAAGCC | CCAGGGACAATGTTTACCACG C |
| Cdh1 (E-Cadherin) | GGTCATCAGTGTGCTCACCTCT | GCTGTTGTGCTCAAGCCTTCA |
| Gsk3b | GAGCCACTGATTACACGTCCAG | CCAACTGATCCACACCACTGT C |
| Vimentin | CGGAAAGTGGAATCCTTGCAG  G | AGCAGTGAGGTCAGGCTTGGA  A |
| Snai1 | TGTCTGCACGACCTGTGGAAAG | CTTCACATCCGAGTGGGTTTG G |
| Cd44 | CGGAACCACAGCCTCCTTTCAA | TGCCATCCGTTCTGAACCACG |
| Hif-1α | CCTGCACTGAATCAAGAGGTTG C | CCATCAGAAGGACTTGCTGGC T |
